# Host relatedness and landscape connectivity shape pathogen spread in a large secretive carnivore

**DOI:** 10.1101/816009

**Authors:** Nicholas M. Fountain-Jones, Simona Kraberger, Roderick Gagne, Daryl R. Trumbo, Patricia Salerno, W. Chris Funk, Kevin Crooks, Roman Biek, Mathew Alldredge, Ken Logan, Guy Baele, Simon Dellicour, Holly B Ernest, Sue VandeWoude, Scott Carver, Meggan E. Craft

## Abstract

Urban expansion can fundamentally alter wildlife movement and gene flow, but how urbanization alters *pathogen* spread is poorly understood. Here we combine high resolution host and viral genomic data with landscape variables to examine the context of viral spread in puma from two contrasting regions: one bounded by the wildland urban interface (WUI) and one unbounded with minimal anthropogenic development. We found landscape variables and host gene flow explained significant amounts of variation of feline immunodeficiency virus (FIV) spread in the WUI, but not in the unbounded region. The most important predictors of viral spread also differed; host spatial proximity, host relatedness, and mountain ranges played a role in FIV spread in the WUI, whereas unpaved roads were more important in the unbounded region. Our research demonstrates how anthropogenic landscapes can alter pathogen spread, providing a more nuanced understanding of host-pathogen relationships to inform disease ecology in free-ranging species.

---

Understanding how pathogens spread through populations remains a fundamental challenge. The extent to which pathogen spread reflects movement patterns of their hosts is enigmatic but important for controlling disease ^1, 2^. If pathogen spread mirrors host gene flow, host genetic structure/differentiation could be a valuable proxy for pathogen spread and be used as a basis to inform disease control ^3, 4^ (e.g., male vampire bat [*Desmodus rotundus*] genetics closely mirrors phylogenetic structure of rabies ^5^). A close relationship between host gene flow and pathogen spread may also be evidence for increased transmission between related conspecifics, and could affect evolutionary pressures on the pathogen, as closely related hosts may be more likely to have similar imune environments ^6^. In contrast, if host gene flow and pathogen spread are decoupled, fine-scale patterns of host movement, for example, may best predict spread and thus inform the strategy employed for disease control ^2, 7^. The patterns of pathogen spread can also be influenced by characteristics of the pathogen itself, where host-specific and directly transmitted pathogens likely have the greatest concordance with host gene flow, relative to multi-host and environmentally transmitted pathogens. Given this range of scenarios, our ability to estimate pathogen spread in heterogeneous landscapes, requires a better understanding of how host relatedness and environmental predictors may drive these processes.

Urbanization is one of the most destructive and large-scale of all anthropogenic landscape fragmentation processes, but how urbanization shapes pathogen spread in particular is still not well understood ^8, 9^. As urban development fragments habitats and introduces barriers (the wildland urban interface, WUI), it can cause reduced host gene flow between populations ^7, 10^, altered animal behaviour (for example, animals becoming more nocturnal to avoid humans ^11, 12^), and changes in feeding ^13^ and movement ^14^ patterns. If these anthropogenic impacts on host behaviour affect transmission dynamics, they may manifest in the demographics of pathogen populations ^15^ (e.g., if transmission events are happening rapidly, the pathogen’s effective population size may be exponentially increasing ^15^). Comparing the factors that shape pathogen spread in populations that are affected by urbanization is often difficult due to a lack of high-resolution data (i.e., coupled host and pathogen genomic data for most individuals) or comparable populations (i.e., well-sampled populations impeded and unimpeded by urbanization). Quantifying how urbanization can affect host gene flow, and how this in turn impacts the transmission dynamics and spread of pathogens, can help address this important research gap.

Here we determine how landscape variables (including those associated with urbanization) and host relatedness affect pathogen spread and transmission in puma (*Puma concolor*). Puma are useful indicators of the effects of urbanization on wildlife as they are sensitive to urban development ^13, 16^ but can persist in areas impacted by urbanization provided sufficient landscape connectivity (e.g., ^17^). As puma foraging, movement, and other behaviours are altered by urban development ^13, 18, 19^, this species offers a valuable case study for how pathogen spread can be effected by urbanization - subject matter that is increasingly important for wildlife conservation and management ^20, 21^. We utilize data we collected from 217 pumas sampled from two geographically distinct regions (approximately 500 km apart): one region bounded by the urban-wildland interface (hereafter the WUI) and the other in a more wild and rural setting relatively unbounded by anthropogenic development (hereafter UB). Our previous work found limited gene flow between pumas from these two regions but similar levels of genetic diversity ^22^. From individuals in both regions, we collated high resolution host genomic and spatial data alongside puma feline immunodeficiency virus (FIV_pco_) sampled from the same individuals. FIV_pco_ is a rapidly evolving retrovirus ^23^ endemic to puma populations, and is thought to be predominantly transmitted horizontally via aggressive encounters ^24^, although vertical transmission has been documented by phylogenetic analyses ^25^. Because FIV_pco_ is essentially apathogenic in puma ^26, 27^, it is an ideal model pathogen to understand transmission dynamics in wild systems without potential confounding effects of disease on behaviour and demography ^25, 28^. We examine what factors impact FIV_pco_ spread using a novel pipeline synthesizing phylodynamic, phylogeographic and landscape genetic techniques (an ecophylogenetic approach ^29^). We employ this pipeline to test for (1) differences in FIV_pco_ demographic histories and transmission dynamics across regions, (2) concordant patterns of host relatedness, viral phylogenetics and spatial distance, and (3) the relative roles of host relatedness and landscape predictors, such as urban development, in shaping the pattern of spread of the virus. We hypothesized that as anthropogenic factors impact puma movement (e.g., ^30^) and gene flow ^22^ at the WUI, that transmission opportunities would be restricted and spatial proximity and host relatedness would be more important in shaping spread in this region.

## Results

Of the 217 individuals we tested we found FIV_pco_ prevalence was higher in the WUI than UB puma (41% vs 59%, see Table S1 for site and population characteristics). We sequenced FIV_pco_ from a total of 46 animals representing most of the infected animals in both populations over a ten-year period. For 43 of the pumas, we obtained both FIV_pco_ sequences (the conserved *pol, ORFA* and *env* genes representing 36% of the FIV genome) and the corresponding puma genomic data (consisting of a data set of 12,444 neutral single nucleotide polymorphisms [SNPs] per individual ^22^).

### Region-specific viral demographic histories

We found not only distinct demongraphic histories in the viruses circulating in the WUI and UB regions, but also differing FIV_pco_ subtypes. Bayesian time-scaled phylogenetic analysis of the FIV_pco_ sequences revealed two co-circulating FIV_pco_ subtypes: FIV_pco_ CO, circulating among pumas in both regions (Fig. 1a) and FIV_pco_ WY, which was only detected in the UB after previously detected in puma in Wyoming (Fig. 1b; see Fig. S1 for a maximum likelihood tree that illustrates the broader phylogenetic context of these two subtypes across North America). Within FIV_pco_ CO, we identified three clades (I, II and III, Fig. 1a) that had contrasting and landscape-specific demographic histories (Fig. S2). Clade I had been circulating predominantly in the WUI since approximately 1995 (95% high posterior density interval (HPD): 1984- 2003), and the effective population size of this clade has been gradually increasing through time (Fig. S2). Clade II showed a similar trajectory in population size (Fig. S2) and was found in puma from both regions, which is potentially indicative of long-distance dispersal of FIV_pco_ (Fig. 1a). Clade III, in contrast, predominantly circulated in the UB, but had a much more distinctive demographic pattern (Fig. S2). We estimated that Clade III began circulating in the WUI in 2001 (95% HPD: 1992-2006) and arrived in the UB in 2006 (95% HPD: 2003-2008), afterwards going through a period of population growth which plateaued around 2012 (Fig. S3). In contrast, we found that FIV_pco_ WY has likely circulated at low prevalence in the UB for over a hundred years (Fig. S3) and had a slower estimated evolutionary rate than FIV_pco_ CO (Table S2).

**Fig. 1.**
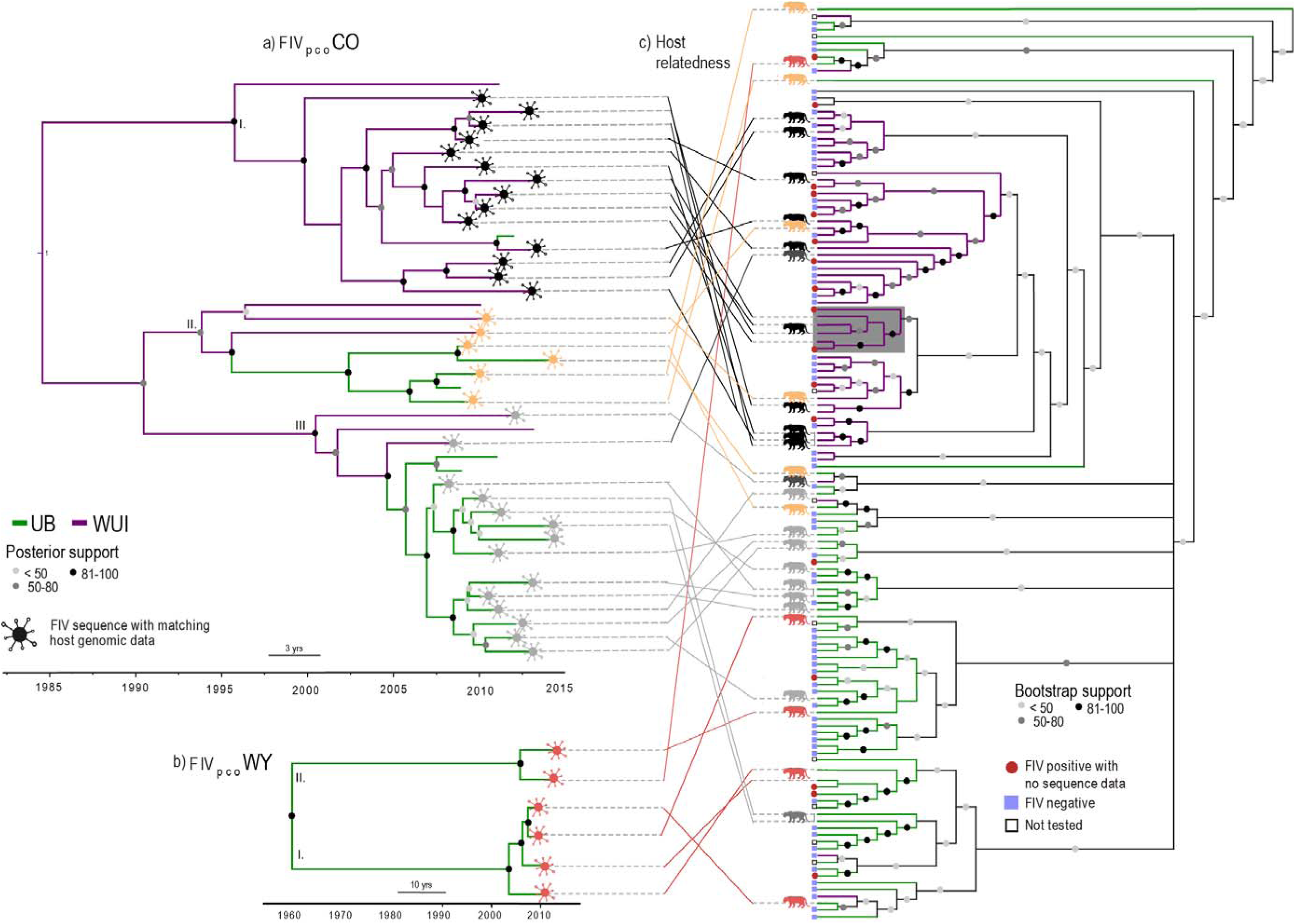
Tanglegram revealing how FIV_pco_ phylogenetic relationships (a & b) overall mapped imprecisely onto the puma relatedness cladogram (c) in our study regions. a) Bayesian time-scaled phylogenetic tree for FIV_pco_ subtype CO found in both the wildland-urban interface (WUI) and unbounded (UB) regions. b) Bayesian time-scaled phylogenetic tree for FIV_pco_ subtype WY which was only found in the UB. I-III represent the different clades identified using tree structure analysis ^31^. Virus branch colours are based on population assignment posterior values from our FIV_pco_ subtype CO discrete trait analysis. c) Host relatedness cladogram constructed using singular value decomposition (SVD) quartets ^32^ based on over 12,000 SNPs from 130 individual puma across both study areas. The grey shaded box encompasses related individuals with phylogenetically similar FIV_pco_ isolates. Virus symbols (from panels [a] & [b]) are coloured based on viral lineage membership. This colour matches the puma infected with that isolate (puma silhouettes [c]) and the lines connecting each isolate to each host in the tanglegram. Tips without virus symbols indicate that there was no matching host genomic data for this FIV_pco_ isolate. Branch colours indicate which region each individual puma and matching virus was sampled from (WUI or UB).

### Divergent patterns of viral and host relatedness across regions

Overall, despite regional fidelity, the FIV_pco_ phylogeny did not map closely onto the puma relatedness cladogram; yet there was some localized evidence for concordance between the two in the WUI region (grey boxes, Fig 1 a/c). For example, four related individuals in the WUI were infected with phylogenetically similar FIV_pco_ (dark grey box, Fig 1 a/c) and were also captured in close spatial proximity to each other (dashed circle, Fig. 2d). In contrast, there was limited evidence of similar patterns in the UB as the most phylogenetically similar FIV_pco_ isolates were sampled across unrelated individuals.

**Fig. 2.**
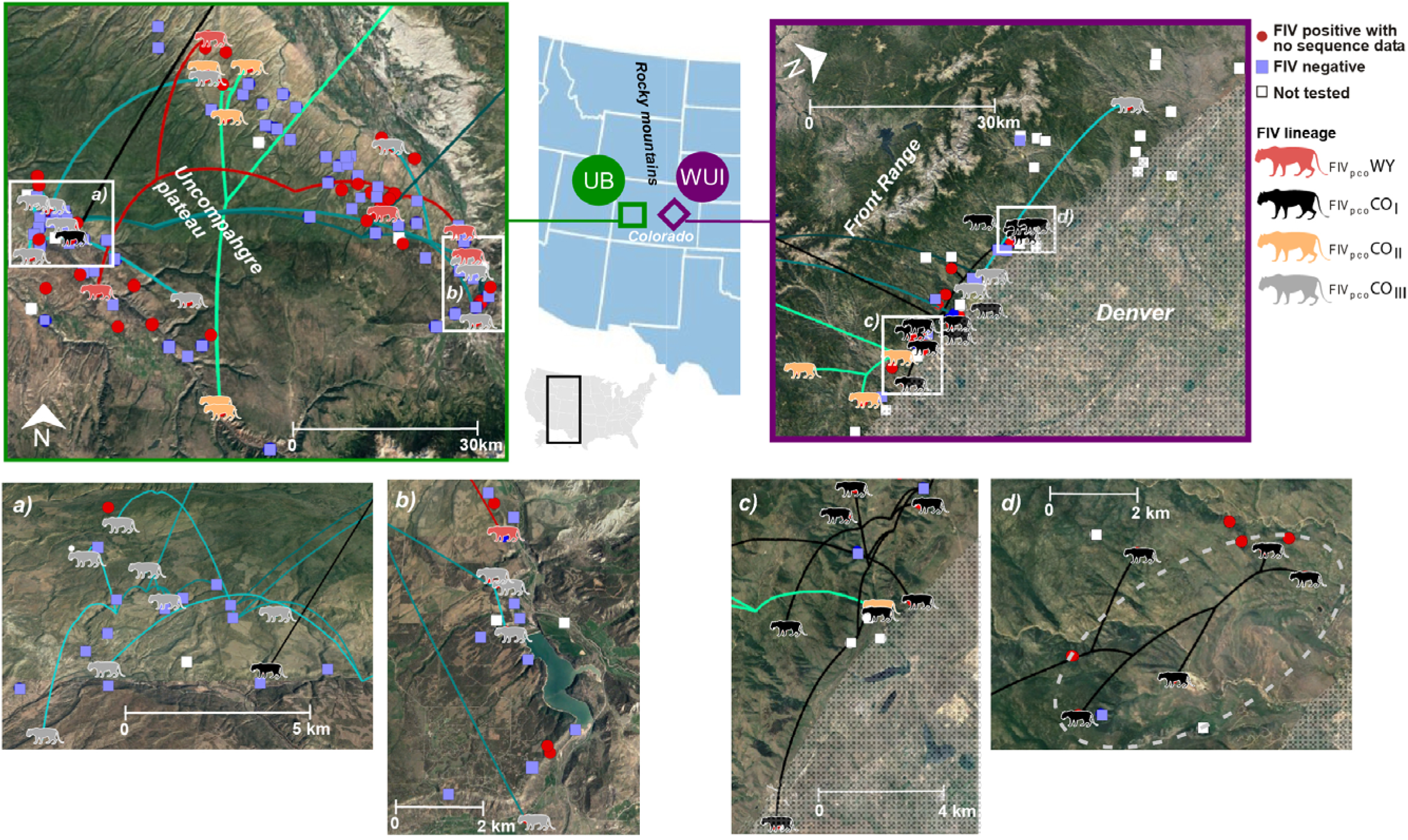
Spatially projected FIV phylogenies showing the configuration of FIV_pco_ spread in the unbounded (UB) and wildland-urban interface (WUI) regions (a-d represent enlarged sections of each area). Tree tips represent the capture locations. Lines indicate FIV_pco_ estimated branch locations, with colours of each line as well as the coloured puma symbols reflecting FIV_pco_ subtype or lineage membership based on tree structure analysis ^31^. The dashed circle indicates the group of individuals in the WUI in which the viral phylogeny mapped onto the host cladogram (grey boxes in Fig. 1). Grey shading shows the extent of the Denver metropolitan areas. Branches coming from outside the study area in the top panels are the branches connecting each region. See Appendix S1a-d for .kml files to recreate this map. FIV_pco_ or host genomic data could not be tested in some individuals (white boxes).

In the UB, a significantly higher proportion of individuals in each ‘neighbourhood’ (i.e., pumas likely to have home range overlap) were qPCR negative for FIV_pco_ than puma in the WUI (Mann-Whitney U Test, *p* = 0.007, Fig. S4). Additionally, in both regions, there was qualitative evidence for spatial structuring with some FIV_pco_ lineages being locally dominant (e.g., CO clade I and III in Fig. 2). However, overall FIV_pco_ spread was a complex mixture of local and longer-distance jumps across both landscapes with uninfected individuals captured at the same time and within 500m of infected individuals in both regions (Fig. 2). Similarly, individuals captured within months of each other at the same location were commonly infected with phylogenetically distinct FIV_pco_ subtypes or clades (Fig. 2).

### Predictors of FIV spread are region-dependent

We employed two complementary techniques to test how host and landscape shaped two components of spread in each region (the overall phylogeographic pattern and lineage dispersal velocity). Strikingly, host and landscape variables explained significant variation of of FIV_pco_ spread for the WUI only. We used generalized dissimilarity models (GDM ^33^) of FIV_pco_ patristic distance (the sum of the lengths of the branches that link two nodes in a tree) to test if landscape and host variables explained variation in the overall viral phylogeographic patterns. In the WUI, our GDM models explained 20% of total model deviance (*p* = 0.028); but only 7% in the unbounded population (*p* = 0.23). Moreover, the most important variables that shaped spread in each case were different (Fig. 3a). In the UB, even though insignificant, FIV_pco_ spread was associated with more roads (i.e., individuals more connected by roads had similar FIV isolates, Fig. 3b). In contrast, the viral spread in the WUI was shaped by spatial proximity coupled with host relatedness and impervious surface. As spatial proximity decreased, so did the FIV_pco_ patristic distance between individuals; neighbouring individuals shared more phylogenetically similar FIV_pco_ isolates (Fig. 3c). Similarly, as host relatedness decreased, so did FIV_pco_ patristic distance (i.e., related individuals were more likely to share phylogenetically similar FIV_pco_ isolates, Fig. 3e). We found a similar positive relationship between FIV_pco_ patristic distance and impervious surface resistance (Fig. 3e). Furthermore, in complement to our relatedness measure, we also included host genetic resistance in our GDM models (see *Methods* for details). Individuals in the WUI with low host genetic resistance values had more similar phylogenetically FIV_pco_ isolates (Fig. 3f). However, this relationship plateaued with dissimilarity values of over 0.05 and was not significant (*p* = 0.38).

**Fig. 3.**
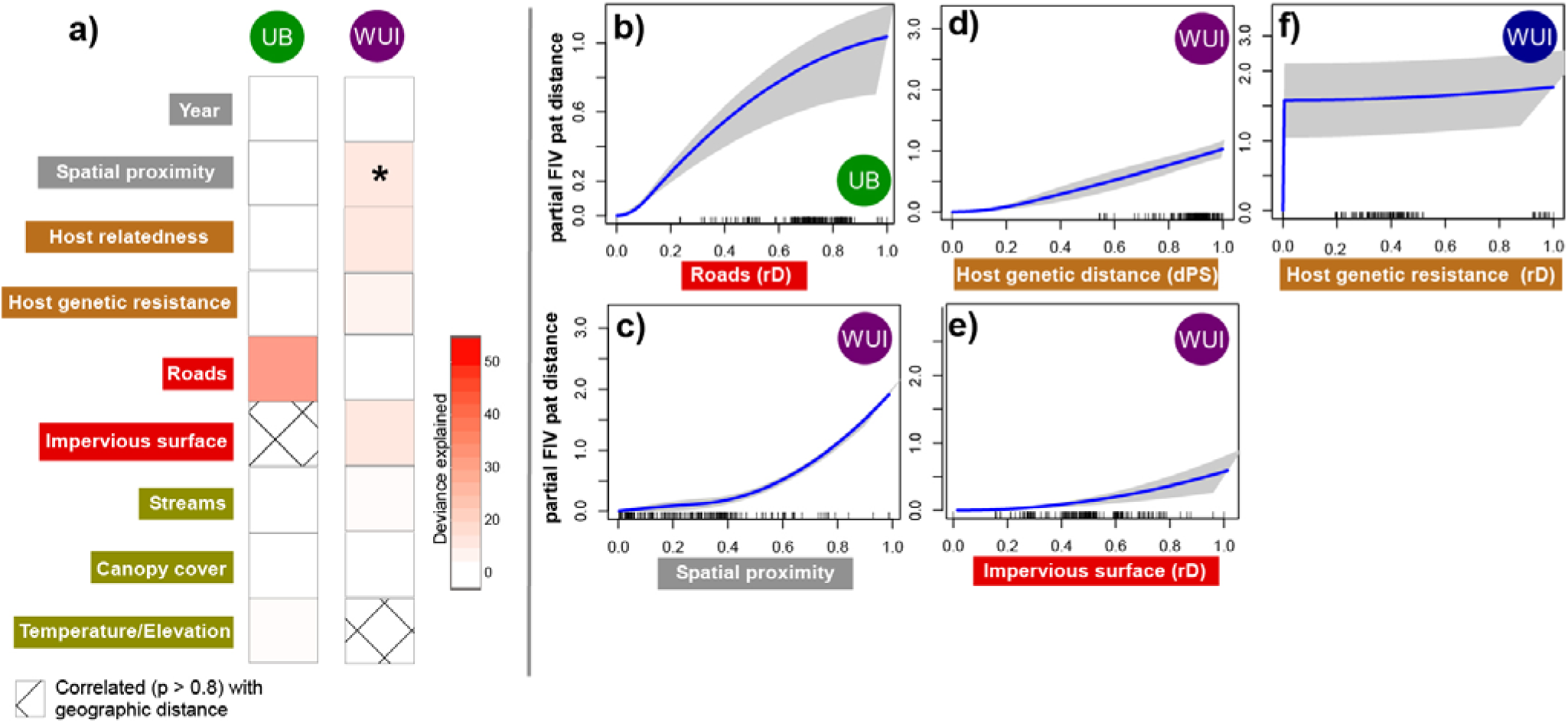
Results from the GDM analyses of FIV_pco_ spread. a) Heat map showing the deviance explained by each predictor in the unbounded (UB) and wildland-urban interface (WUI) *: *p* value 0.01-0.05. Predictor labels in grey boxes = spatiotemporal predictors, predictors in orange = host genetic measures, predictors in red = associated with anthropogenic development, predictors in olive green = natural landscape features. Right panel shows GDM-fitted I-splines showing the relationship between partial FIV_pco_ patristic distance for the variables selected in the UB final model (b, see *Methods*) and WUI (c-f) in order of importance. Pat distance = patristic distance. Lines across the x-axis represent a rug plot showing the distribution of the data and the grey surrounding the blue line (the fit) indicates 95% confidence intervals. rD: Resistance distance. dps: Proportion of shared alleles.

We also found that there were region-specific impacts of landscape on FIV_pco_ lineage dispersal velocity. Our analysis revealed that elevation tended to act as a conductance factor increasing the dispersal velocity of FIV_pco_ lineages in the WUI, whereas none of predictors we measured had any substantial effect on lineage dispersal velocity in the UB (positive *Q* distribution and associated Bayes factor support >3 ^34^; see Fig. S5 and Table S3 for a list of tested landscape factors). Furthermore, warmer minimum temperatures (as measured in the coolest month) tended to act as a resistance factor decreasing lineage dispersal velocity in the same region. Taken together, these results indicate that in the WUI, FIV_pco_ tended to spread faster through higher elevation areas less suitable for puma habitat. In the WUI, the areas of higher elevation tended to be away from the urban edge.

## Discussion

### The connections between host and pathogen at the urban edge

Our multifaceted approach linking landscape, host genomic, and pathogen genomic data uncovered unique landscape-specific effects on FIV_pco_ spread, indicative of altered epidemiological dynamics associated with urban landscape structure. We found that spatial proximity coupled with host relatedness and impervious surface was positively associated with viral spread, but only in the WUI (Wildland-Urban Interface). These landscape factors mirrored the factors shaping host gene flow ^22^ providing evidence that host movement and viral spread were more intimately related in the WUI than in the UB (Unbounded) region. However, whilst there was some evidence for concordance between the FIV_pco_ phylogeny and host cladogram in the WUI, for the most part they did not map precisely onto each other in either region (Fig. 1) indicating that transmission was not only happening just between related individuals. There was little evidence of FIV_pco_/host concordance in the UB where host relatedness did not shape phylogeography and entirely different sets of predictors shaped host gene flow (spatial proximity and tree cover ^22^). These opposing patterns between host gene flow and viral spread could reflect regional differences in transmission. One potential scenario is that transmission between neighbouring related conspecifics may be more likely in the WUI due to altered puma movement and dispersal patterns ^18, 19^ and foraging behaviours ^13^. The urban development that impacts the WUI is linear (i.e., where the great plains meet the Front Range of the Rocky Mountains). This linear development restricts juvenile dispersal ^19^ and could lead to more opportunities for transmission among more related conspecifics (i.e., individuals establish home ranges close to their parents). Our previous work supports this hypothesis as we demonstrated that family units were more clustered in space, where puma genetic distance per kilometre and sub-structure was greater in the WUI compared to the UB ^22^. There has not been an extensive evaluation of relatedness in neighbouring home-range females, but female matrilines (groups of maternally-related females) are known to occur ^35, 36^. Furthermore, we found evidence that infection was more clustered in space in the WUI compared to the unbounded one, supporting the idea that spatial proximity increases transmission risk between related individuals in the WUI. This shift in transmission risk may reduce evolutionary pressure on the virus due to factors such as similarity in immune profile ^6^.

### Host gene flow and viral phylogeography uncoupled in the unbounded region

Roads, common but mostly unpaved in this UB region, were a modest predictor of spread in the UB. Radiotelemetry has shown that puma often move using unpaved roads^37^ and rapid viral evolution potentially allowed us to detect this modest effect. There was a smaller impact of roads on host gene flow ^22^ which supports the idea that FIV phylogenetics can capture more contemporary movement patterns impossible to detect using host genetics alone ^7, 38, 39^. When host gene flow and viral spread are decoupled, as was the case in the UB, the rapid accumulation of viral mutations may conversely obscure historical trends in connectivity ^40^. This could explain why we detected no effect of tree cover on FIV_pco_ spread even though puma are known to have a preference for tree cover to disperse and hunt ^41, 42^. The altered epidemiological history of the clade, dominant in the UB compared to all other detected clades, may reflect or be a consequence of relatively unrestricted spread in the UB (Fig. S3, FIV_pco_ CO clade I). We postulate that recent arrival of FIV_pco_ CO clade I in the UB and signature of rapid expansion ^15^ across the Uncompahgre Plateau (Fig. 2, Appendix S1) may only have been possible in a region where viral spread itself was unbounded. Further work is needed to assess the temporal dynamics of FIV_pco_ in the UB. The high elevation Uncompahgre Plateau (averaging 2900 m a.s.l, Fig. 2) did not shape viral spread, even though puma are known to have a preference for not dispersing across high altitude divides^41^. In contrast, we found that higher altitude areas increased dispersal velocity in the WUI. As most human activity in the Front Range (Fig. 2) is in lower altitude areas, it is plausible that increased viral velocity in higher altitude areas is a product of avoidance of the human ‘super predator’ ^13, 43^. FIV transmission events have been shown to be more likely to occur further from the urban edge in bobcat populations ^22^ and the same may apply for pumas in the WUI region here. Velocity may also be faster through higher elevations in the Front Range as puma are more likely to rapidly move through unsuitable habitat. Unsuitable habitat has also been demonstrated to increase the velocity of rabies lineages in dogs ^44^.

### Disease management implications

Our findings have pathogen and host management implications, as we demonstrated that spatial proximity and host relatedness may be relevant predictors of pathogen spread in regions impacted by urban development. This may mean that the difficult task of disease control in a large apex predator (such as vaccinating against feline leukaemia virus in a puma population ^21^) may be more tractable in bounded populations. Because juveniles set up home ranges near their parents, dispersal events important for pathogen jumps in the landscape are more constrained. Further, as host gene flow and viral spread were tightly linked in the WUI, targeting individuals where gene flow was less constrained by impervious surface (further from the urban areas) may also reduce spread. We acknowledge that in this study we did not sample all pumas in the system nor between the two sampled regions, and that FIV_pco_ could not always be sequenced. This could mean that we missed, for example, some FIV_pco_ lineages that could have altered our inference about FIV_pco_ patterns. Nonetheless, this did not compromise our ability to gain complementary insights into the drivers of host connectivity by combining high-resolution host and pathogen genomic data, which would have been impossible to detect with either host or pathogen data alone. Our work provides a valuable case study of how landscape context and host relatedness is likely to be important in disease management plans. As urban landscapes continue to expand, improving our understanding of how heterogeneous landscapes and host relatedness alter pathogen transmission will be increasingly important.

## Methods

### Samples

Puma blood and tissue samples were collected from 103 individuals (48 males, 55 females) between 2005-2014 in the UB and 110 individuals (43 males, 54 females, 12 undetermined) between 2003-2015 from the WUI as part of monitoring efforts by Colorado Parks and Wildlife in the Rocky Mountain Range of Colorado, USA ^19, 45^. This sampling effort is likely to represent a large proportion of the resident puma present in both regions during the sampling period ^19^. See Table S1 for further details on the samples.

### FIV detection and sequencing

Total DNA was extracted from 50µl whole blood samples using the QIAGEN DNeasy Blood & Tissue extraction kit (Qiagen, Inc., Valencia, CA) with an extended incubation period of two hours or from 200 µl whole blood samples using a phenol chloroform extraction as per^46^. Isolated DNA was quantified using a QuBit 2.0 fluorometer (ThermoFisherScientific). DNA from individual puma were screened for the presence of FIV_pco_ provirus using a specific qPCR assay as described by ^47^. Full *ORFA* and *pol* gene regions were isolated from those samples identified as qPCR positive using a nested PCR protocol. See Text S1 for the sequencing protocol and Table S4 for primer details. While we also sequenced the *env* gene, our assessment of the temporal signal (see next section) indicated many discrepancies in the data regarding the use of a molecular (strict or relaxed) to analyse these data. Resulting genetic sequences with chromatograms were checked, assembled, trimmed and aligned using the MUSCLE algorithm ^48^ using Geneious 7.0.6. Our *Pol* and *ORFA* datasets (GenBank accession: MN563193 - MN563239) were compiled together with those sequences available in the public database Genbank, previously isolated from across the USA ^49^ (Fig. S1). Recombination was detected using RDP software V4 ^50^; parameters were set at default with linear topology. Events were determined as true if supported by three or more methods with *p* values <10e^-3^ combined with phylogenetic support. Recombination-free datasets were used for all downstream phylogenetic analyses.

### Viral Phylogenetics

To examine the broad placement of the FIV_pco_ isolates sampled during this study, a maximum-likelihood tree was constructed for the FIV_pco_ dataset comprised of all isolates recovered in the USA using PhyML ^51^with the TN93+G+I model and aLRT branch support, branches with <80 support were collapsed using TreeGraph2 ^52^.

We used TempEst to assess data quality control of our generated data through root-to-tip regression and observed largely varying deviations from the regression line for many of the sequenced *env* genes. These deviations precluded the use of strict or relaxed molecular clock models to analyse the data, and we hence decided to move forward with the *pol* and *ORFA* sequence data. We used BEAST 1.10 ^53^ with BEAGLE 3.1 ^54^ to perform both discrete and continuous phylogeographic analyses ^55^ based on the concatenated *pol* and *OFRFA* sequences from each subtype using HKY substitution model (found most suitable for this smaller number of sequences using smart model selection ^56^). For FIV_pco_ CO, we performed Bayesian model selection on various model combinations comprising a strict molecular clock and an uncorrelated relaxed clock model with an underlying lognormal distribution, as well as three different coalescent models: a constant population size model, an exponential growth model and a non-parametric Bayesian skygrid model ^57^. We also tested three relaxed random walk (RRW) models of continuous diffusion for the phylogeographic analyses and included each population (WUI and UB) as a discrete trait. All Bayesian model selection experiments were performed by (log) marginal likelihood estimation using path sampling and stepping-stone sampling ^58–61^ (see Table S5). Multiple replicates were run with different starting seeds to ensure convergence. Based on the outcome of the Bayesian model selection procedure, we presented the results obtained for the relaxed molecular clock models, the exponential population size coalescent model and the Cauchy RRW model. For FIV_pco_ WY, as there were only 6 sequences, we applied a different approach as there was not enough data to get stable results for complicated evolutionary models. In this case, we assumed a strict clock and constant population size and set the root prior to a uniform distribution (0, 300) reflecting our expectation that this subtype had been circulating for no more than 300 years. For both subtypes, duplicate MCMC chains were run for 200 million generations, with trees and parameters sampled every 20,000 steps. We used the program Tracer version 1.7 ^62^ to examine ESS values (with parameter estimates accepted if the ESS was > 200) and obtain HPD intervals for estimated parameters.

To identify hidden population structure in our time-scaled phylogenies, we applied the ‘tree structure’ R package ^31^ using the default values. This analytical routine compares discrepancies between observed and idealised genealogies to identify clades under differing epidemiological or demographic processes ^31^. As we did not have enough sequences to find meaningful structure in FIV_pco_ WY, we limited this analysis to FIV_pco_ CO. Any clades identified as significantly departing from the idealised geneology were further interrogated using the R package ‘phylodyn’^63^ to estimate effective population size through time. This nonparametric Bayesian approach uses integrated nested Laplace approximation (INLA) to estimate effective population size efficiently, whilst accounting for preferential sampling by modelling the sampling times as a Poisson process (see ^64^).

### Host genomic data

We successfully genotyped 130 pumas (76 individuals from the UB and 54 individuals from the WUI) using a ddRADseq approach (see ^22^ for sequence and bioinformatics details). From these genomic data, we quantified individual relatedness using the inverse proportion of shared alleles (Dps ^65^) and used the resultant pair-wise distance matrix in the downstream analyses. We were unsuccessful in getting ddRAD data for seven individuals for which corresponding viral genomic data were available (see Fig. 1) and for these individuals we used the mean population relatedness value. Removing these individuals from the analyses did not qualitatively alter the results. Furthermore, we used the complete Dps dataset from all individuals and interpolated this distance across each landscape using a kriging approach. We converted this interpolated surface into a resistance raster in R and calculated resistance distances between each individual with FIV_pco_ sequence data using a Circuitscape model, which uses circuit theory to accommodate uncertainty in the route taken ^66^. See https://github.com/nfj1380/ColoradoPumaFIVproject for details.

We estimated a coalescent-based phylogeny with bootstrap support values using SVD quartets ^67^ as implemented in PAUP ^68^. SVD quartets, which is currently considered the most robust and computationally efficient SNP-based phylogenetic estimation method^32^ and uses a site-based approach (considering each SNP has an independent genealogy) to estimate combinations of four-taxon relationships and heuristically summarise the resulting trees into a species tree phylogeny^32, 67^. We evaluated a maximum of 100,000 random quartets using the QFM quartet assembly method and the multispecies coalescent tree model for generating the topology, and performed 100 bootstrap replicates for assessing topological branch support for all terminal taxa.

### Landscape Data

We collected Geographic Information Systems (GIS)-based landscape data that we hypothesized would be important for puma relatedness in Colorado (see Table S3 for more detail on GIS data sources and ecological justification for each landscape variable). This included percent impervious surface (e.g., roads, buildings, etc.), land cover (forested, open- natural, and human developed as sub-rasters), percentage tree canopy cover, vegetation density, rivers/streams, roads, minimum temperature of the coldest month, annual precipitation, topographic roughness, and elevation. Resistance surfaces were created from this landscape data using the Reclassify and Raster Calculator tools in ArcGIS v. 10.1. We calculated resistance distances between individuals with FIV_pco_ sequence data also using Circuitscape path model.

We calculated the proportion of uninfected to infected individuals within a 5 km buffer (estimated average distance between individuals ^69^) of each FIV_pco_ positive individual using ‘summarize within’ tool also in ArcGIS v. 10.1. Estimates from each population were compared using a Mann-Whitney U Test.

### Impact of host relatedness and landscape resistance on viral spread

We used generalized dissimilarity modelling (GDM) ^33^ to quantify if host relatedness and landscape shaped FIV_pco_ spread for FIV_pco_ CO. GDM is a flexible nonlinear regression approach that fits monotonic I-spine functions to pair-wise matrix data ^33^ to describe the rate and magnitude of, in this case, FIV_pco_ phylogenetic change. We first calculated FIV_pco_ patristic distance (using the maximum clade credibility tree) and converted this distance metric into a dissimilarity measure 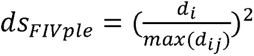 where *d* is pairwise distance. The host Dps matrix and all of the landscape resistance matrices were converted to dissimilarities the same way. Specifically, GDM uses generalized linear models (GLMs) to model FIV_pco_ patristic distance in the form of:

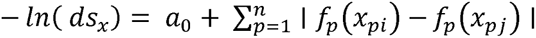

where *i* and *j* are individual puma, *a*_0_ is the intercept, *p* is the number of covariates and *f_p_(x)* have I-spline transformed versions of the predictors (see ^33, 70^ for further details). We used a backward elimination model selection approach and permutation tests (n = 99) to test for significance ^33, 71^. The model with the highest deviance (±2% deviance explained) with the smallest number of predictors was reported. We performed GDM using the same predictor sets as ^22^ for each region but in our reduced dataset (i.e., with individuals with FIV_pco_ data), temperature and elevation were strongly correlated with space (Mantel *r* = 0.93). We tested different values of K (see below) and treated each as resistance or conductance surfaces yet this made no difference to the GDM models (we present results from resistance surfaces only with K=100). Analysis of FIV_pco_ WY was not possible given the small sample size.

### Impact of environmental factors on viral dispersal velocity

The analysis of the impact of environmental factors and host genetic differentiation on the dispersal velocity of viral lineages was performed using R functions of the package “seraphim” ^6^ (see ^65, 66^ for a similar workflow). In this analysis, each environmental factor, as well as the interpolated host genetic distance surface, was described by a raster that defines its spatial heterogeneity and that was used to compute an environmental distance for each branch in the phylogeny using two different path models: (i) the least-cost path model, which uses a least-cost algorithm to determine the route taken between the starting and ending points ^75^, and (ii) the Circuitscape path model. Here, we investigated the impact of the environmental rasters listed in Table S3 as well as the resistance raster generated from host genetic distance interpolation. We generated distinct land cover rasters from the original categorical land cover raster (resolution = 0.5 arcmin) by creating lower resolution rasters (2 arcmin) whose cell values equalled the number of occurrences of each land cover category within the 2 arcmin cells ^73^. For each considered environmental factor, several distinct rasters were also generated by transforming original raster cell values with the following formula: v_t_ = 1 + *k*\*(v_o_/v_max_), where v_t_ and v_o_ are the transformed and original raster cell values, and v_max_ the maximum raster cell value recorded in the raster. The rescaling parameter *k* here allowed the definition and testing of different strengths of raster cell conductance or resistance, relative to the conductance/resistance of a cell with a minimum value set to “1”. For each environmental factor, we tested three different values for k (i.e. 10, 100 and 1000). Finally, all these rasters were tested as potential conductance factors (i.e., factors facilitating movement) and as possible resistance factors (i.e., factors impeding movement). The statistic *Q* was used to estimate the correlations between phylogenetic branch duration and environmental distances. *Q* is defined as the difference between two coefficients of determination (R^2^): (i) R^2^ obtained when branch durations are regressed against environmental distances computed on the environmental raster, and (ii) R^2^ obtained when branch durations are regressed against environmental distances computed on a null raster, i.e., an environmental raster with a value of “1” assigned to all the cells. For positive distributions of estimated *Q* values (i.e., with at least 90% of positive values), statistical support was then evaluated against a null distribution generated by a randomization procedure and formalized as an approximated Bayes factor (BF) support ^76^. To account for the uncertainty related to the Bayesian inference, this analysis was based on 1,000 trees sampled from the post-burn-in posterior distribution inferred using the continuous phylogeographic model. We performed two distinct analyses, one per region, gathering all phylogenetic branches occurring on each study area.

## Supporting information

Supps

