## Supplementary material for "Host relatedness and landscape connectivity shape pathogen spread in a large secretive carnivore": Supps

### Supplementary materials

**Table S1.** Summary of puma samples used in this study, pathogen data and population level host genetic diversity estimates and study area size.

| Population | #Puma sampled | % qPCR^§^ tested± | FIV_pco_ prevalence (%) | % sequenced^ѱ^ | % with host ddRADseq data | Host genetic diversity* | Geographic size of areas sampled (km^2^) |
| --- | --- | --- | --- | --- | --- | --- | --- |
| Unbounded | 103 | 93 | 41 | 70 | 78 | 1.93 | 11889 |
| WUI | 110 | 65 | 59 | 68 | 80 | 1.89 | 11958 |

§: Quantitative polymerase chain reaction. ± Some individuals were not tested due to insufficient blood volume. ѱ: Both full length *pol*, *ORFA* and *env* genes sequenced.*: allelic richness from (1).

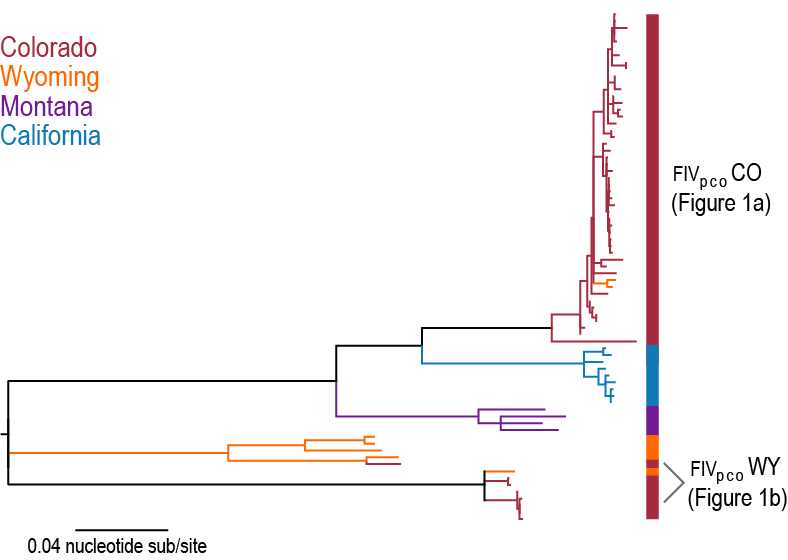

**Fig. S1.** Maximum likelihood tree illustrating the phylogenetic context of both subtypes found in our Colorado population compared to isolates in puma across the western U.S (GenBank accession numbers EF455603 - EF455612, KF906175 - KF906194 and MN563193 - MN563239).

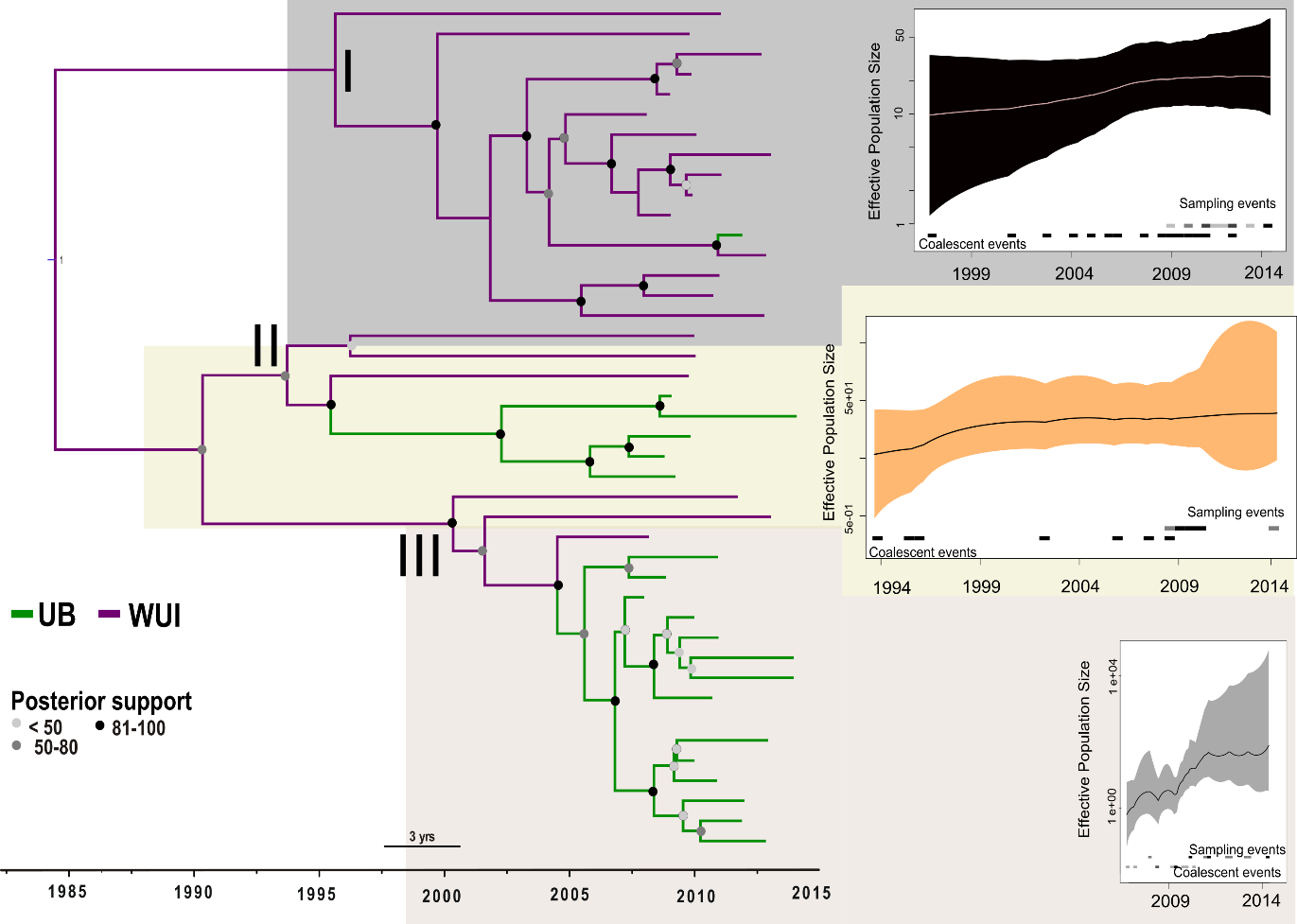
**Fig. S2.** Estimates of effective population size through time for each significantly different FIV_pco_ CO lineage (lineage I, II & 3, *p* = 0.05) based on structure analysis of our Bayesian time-scaled phylogeny (see *Methods* in the main text). The colour of the virus symbols and 95% high posterior density (HPD) intervals reflect each lineage. Edges are coloured based on the puma population the isolate was sampled from (UB: unbounded, WUI: wildland urban interface). Virus branch colours are based on population assignment posterior values from our FIV_pco_ subtype CO discrete trait analysis (see *Methods: viral phylogenetics*). Sampling events refer to dates when samples were collected from puma and coalescent events are estimated mean dates when branches coalesced.

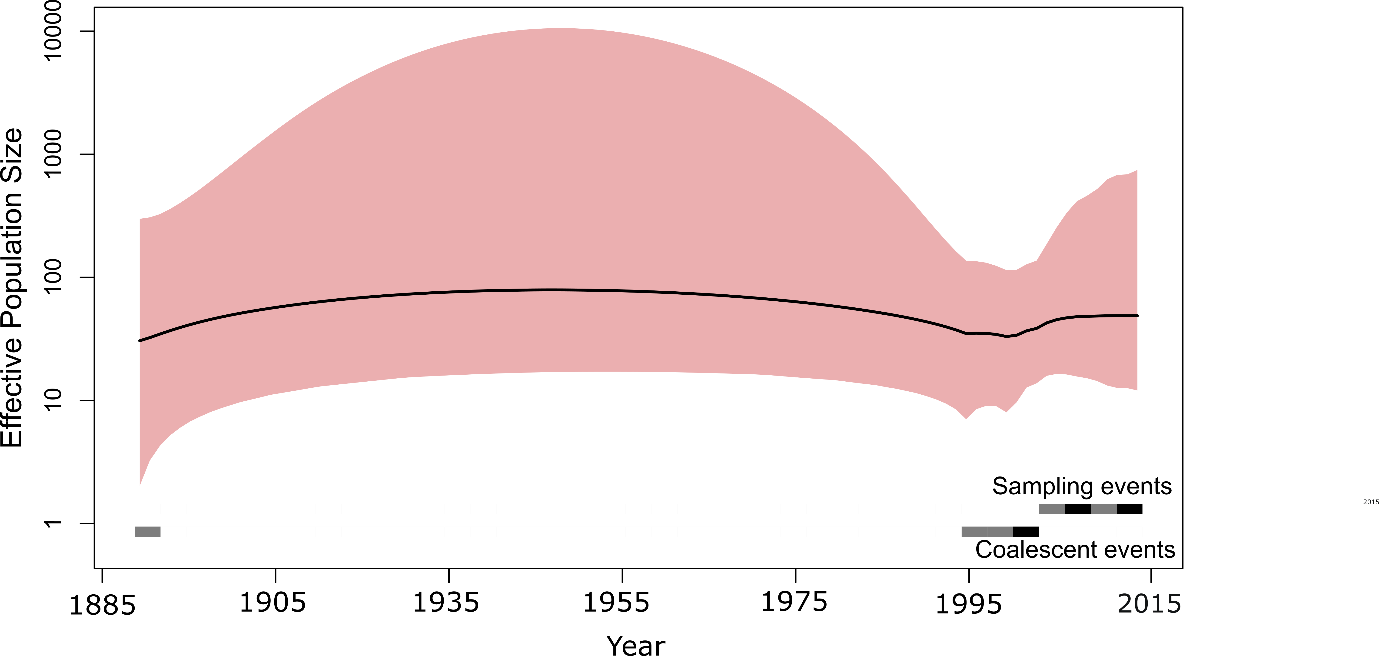

**Fig. S3.** Estimates of effective population size through time for FIV_pco_ WY (95% high posterior density interval in red). Sampling events refer to dates when samples were collected from puma and coalescent events are estimated mean dates when branches coalesced.

**Table S2.** Subtype evolutionary rates.

| **Subtype** | **Evolutionary rate (**substitutions per site/yr) | **95% HPD*** |
| --- | --- | --- |
| FIV_pco_ WY | 1.04e ^-4^ | 2.64e ^-5^, 2.30e^-4^ |
| FIV_pco_ CO | 4.25e^-4^ | 2.48e ^-4^, 6.04e^-4^ |

* highest posterior density.

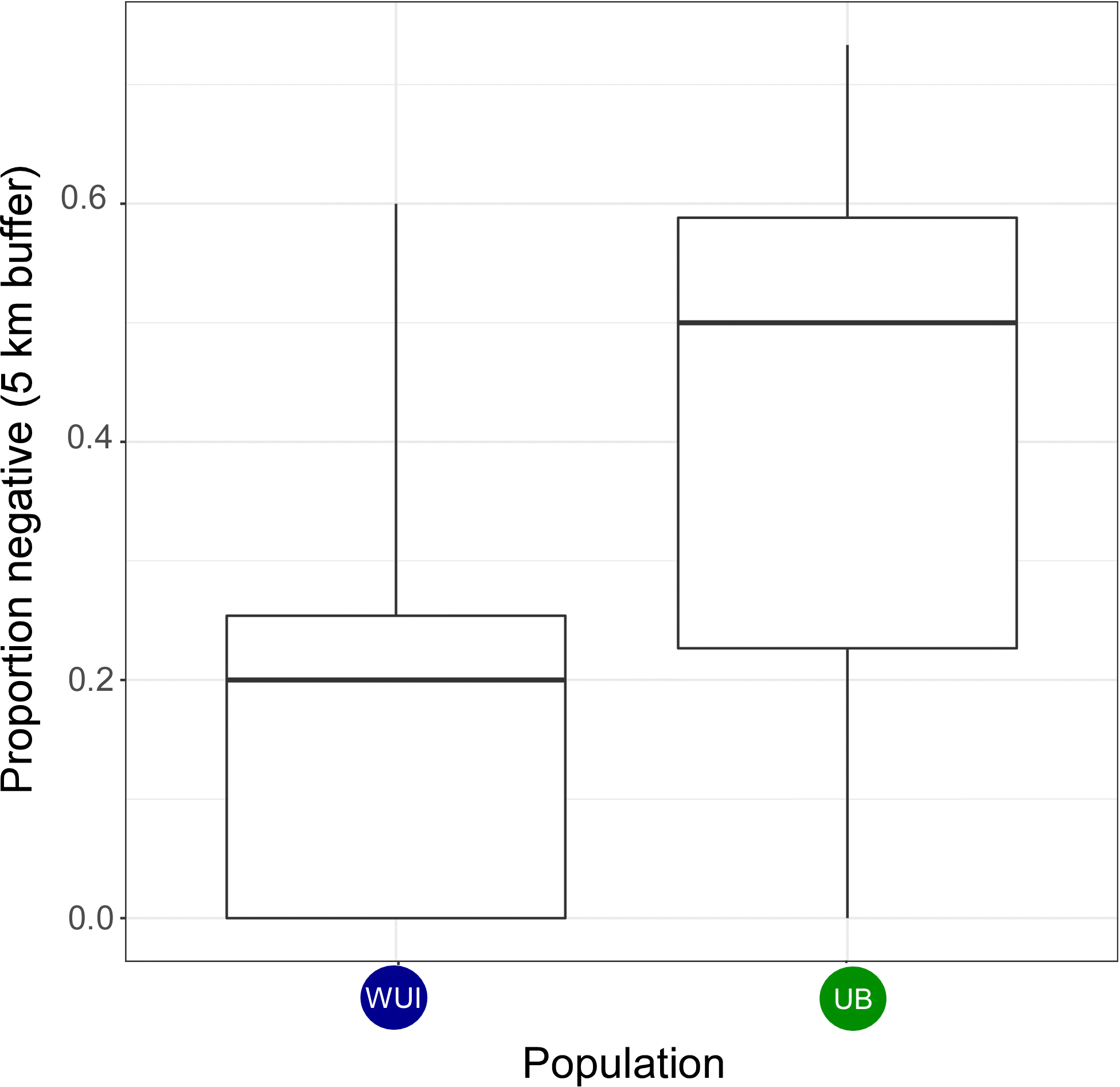

**Fig. S4.** Boxplot showing the proportion of individuals FIV_pco_ negative in a 5 km buffer around a positive individual in each population (WUI: wildland urban interface, UB: unbounded).

**Table S3.** Environmental variables used for landscape genomic analyses, source of the environmental data, and ecological justification for each variable (1)

| **Category** | **Landscape variable** | **Code** | **Description** | **Data source and resolution** | **Calculation** | **Ecological justification** |
| --- | --- | --- | --- | --- | --- | --- |
| Distance | Isolation by Euclidean distance (null model) | Geographic proximity | Euclidean distance between individuals | National Elevation Dataset (lta.cr.usgs.gov/ned) National Map Tool (viewer.nationalmap.gov),  30 meter | Geospatial Modelling Environment (spatialecology.com/gme), ArcGIS 3D Analyst, Circuitscape | Null model of isolation by straight-line distance over variable topography (2). |
| Land cover | Land cover: forested, open-natural, and developed | Land cover | Multiple land cover categories collapsed into 3 costs of movement: forested (lowest), open natural areas (medium), and developed (highest) | National Land Cover Database (mrlc.gov/nlcd2011.php; Homer et al. 2011),  30 meter | ArcGIS Spatial Analyst | Forested habitats provide the most cover for hunting and dispersal, open natural areas are intermediate, and developed areas are the least suitable habitat for dispersal (3, 4). |
|  | Percent impervious surface | Impervious | Percentage of impervious surface | National Land Cover Database (mrlc.gov/nlcd2011.php; Homer et al. 2011),  30 meter | ArcGIS Spatial Analyst | Human development results in increased noise, lights, and hunter access, limiting dispersal (5–7) |
|  | Road corridors | Roads | Roads, with 50 meter buffers on each side | Colorado Department of Transportation (dtdapps.coloradodot.info/otis),  30 meter | ArcGIS Analysis Tools, Spatial Analyst | Roads increase mortality, noise, lights, and hunter access, limiting dispersal (5, 7, 8). |
|  | River and stream riparian corridors | Riparian | River and stream riparian corridors, with 50 meter buffers on each side | National Hydrography Dataset (nhd.usgs.gov),  30 meter | ArcGIS Analysis Tools, Spatial Analyst | River and stream riparian corridors provide vegetative and topographical cover for dispersal, as well as water sources attracting prey species (9, 10) |
| Vegetation | Percent tree canopy cover | Tree cover | Percentage of tree canopy cover | National Land Cover Database (mrlc.gov/nlcd2011.php; Homer et al. 2011),  30 meter | ArcGIS Spatial Analyst | Low tree canopy limits cover for ambush predation and concealment, restricting dispersal (11–13). |
|  | Enhanced vegetation index | Veg. density | Density of vegetation calculated from chlorophyll reflectance in visual and near-infrared spectra | Moderate Resolution Imaging Spectroradiometer (modis.gsfc.nasa.gov),  250 meter | ArcGIS Spatial Analyst | Low vegetation density limits cover for ambush predation and concealment, restricting dispersal (10, 12, 13). |
| Climate | Minimum temperature of the coldest month | Min. temp. | Mean annual minimum temperature of the coldest month (°C) calculated from 1970-2000 weather station data, interpolated between stations | Global Climate Data (worldclim.org/bioclim; Hijmans et al. 2005),  1 kilometer | ArcGIS Spatial Analyst | Lowest minimum temperatures, found at high elevation mountain ridgelines restrict hunting, breeding, and dispersal (14). |
|  | Mean annual precipitation | Ann. precip. | Mean annual precipitation accumulation (mm) calculated from 1970-2000 weather station data, interpolated between stations | Global Climate Data (worldclim.org/bioclim; Hijmans et al. 2005),  1 kilometer | ArcGIS Spatial Analyst | Dry habitats with low precipitation accumulation limit prey species for hunting and vegetative cover, restricting dispersal (11, 15). |
| Topography | Topographic roughness | Topo. rough. | Topographic complexity based on variance in elevation within a moving window | National Elevation Dataset (lta.cr.usgs.gov/ned) National Map Tool (viewer.nationalmap.gov),  30 meter | Geomorphometric and Gradient Metric Toolbox (Cushman et al. 2010), ArcGIS Spatial Analyst | Steep, topographically-complex canyons and mountain slopes provide cover for hunting and dispersal (9, 14). |
|  | Elevation | Elevation | Elevation calculated from digital elevation models. | National Elevation Dataset (lta.cr.usgs.gov/ned) National Map Tool (viewer.nationalmap.gov),  30 meter | ArcGIS Spatial Analyst | Higher elevation, forested habitats are better for hunting, breeding, and dispersal than lower elevation, prairie and high desert habitats. |

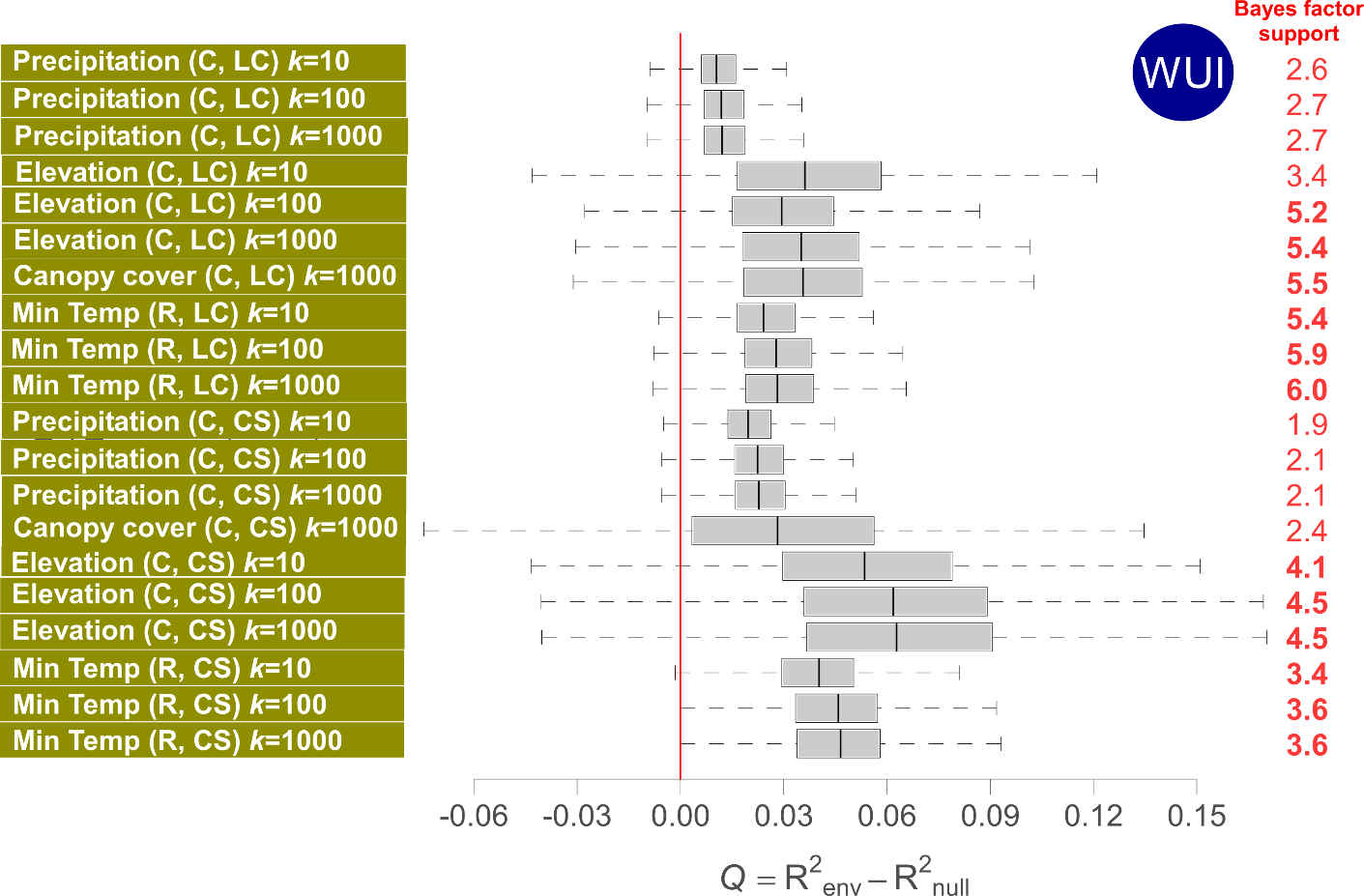

**Fig. S5.** Analysis of the impact of environmental factors on the dispersal velocity of viral lineages. The figure reports the estimated *Q* distributions and associated approximated Bayes factor (BF) support for the environmental factors associated with at least 90% of positive *Q* values. Here, we report results for the WUI population (CO only) as there were no factors with both positive *Q* distributions [p(*Q*>0) > 0.9)] for the unbounded population. “C” and “R” indicate if the considered environmental variable was considered as a conductance (“C”) or resistance factor (“R”), and *k* is the rescaling parameter used to transform the initial raster. “LC” and “CS” indicate if the environmental distances were computed using the least-cost (“LC”) or Circuitscape (“CS”) path model. Following the scale of interpretation of BF values defined by (16), BF values higher than 3 can be considered as “positive” support and values higher than 20 can be considered “strong” support. Approximated BF values higher than 3 are highlighted in bold.

**Text S1.**

*Sequencing Protocol*

Master mix PCR reactions for round 1 - 5 µl Kapa Hotstart hifi polymerase (Kapa biosystems, USA), 2 µl H_2_O, 1 µl of both the forward and reverse primer (see below) and 2 µl of DNA., and in round 2 - 10 µl Kapa Hotstart hifi polymerase (Kapa biosystems, USA), 6 µl H_2_O, 1 µl of both the forward and reverse primer (see below) and 2 µl of R1 PCR product. The PCR cycling conditions were run according to manufacturer’s specifications with an annealing temperature at 60 ºC with reactions run on a C1000 Touch Thermal Cycler (BioRad). Primer details can be found in Table S2 below. PCR products were run on a 0.7 % agarose gel using gel electrophoresis and bands of the correct size excised from the gel. Excised DNA fragments were purified using a MEGAquick-spin™ Total Fragment DNA Purification Kit (iNtRON Biotechnology, Korea). These purified fragments were ligated into pJET 1.2 blunt vector using the CloneJET PCR Cloning Kit (Thermo Fisher Scientific) and cloned using heat shock method with XL1-Blue *E. coli* competent cells (Agilent). Plasmids containing fragments of interest using DNA-spin Plasmid Purification Kit (iNtRON Biotechnology, Korea) and Sanger sequenced using primer walking at Quintarabio (San Francisco, CA).

**Table S4.** List of primers used to sequence FIV_pco_. The majority of sequences were obtained using primers starting with PLVB_[pol/env]. For several poorly amplifying pol sequences we developed an additional round two primer PLVB_PolNewF R2. FIV_pco_ subtype WY *env* was amplified using PLVBWY primers.

| **Name** | **Gene** | **Primer sequence** |
| --- | --- | --- |
| *Round 1* PLVB-PolF R1 | pol | GAATATGTTSGCKCAAGCYTTACAAC |
| PLVB_PolR R1 | pol | GTGRAAGGACCAAAGAATTCCTTCTAC |
| *Round 2* PLVB-PolF R2 | pol | GAATATGTTSGCKCAAGCYTTACAAC |
| PLVB_PolR R2 | pol | CATRTTTCTTTTACTACTCTGYACYCTG |
| PLVB_PolNewF R2 | pol | GCACAAGCGTAACAACAGGT |

**Table S5.** Marginal-likelihood estimates for the candidate models inferred using path-sampling (PS), stepping-stone sampling (SS) for FIV_pco_CO. The model in italics was the model used for subsequent analysis.

| **Model #** | **Relaxed-clock** | **Tree coalescent** | **Continuous trait model** | **SS** | **PS** |
| --- | --- | --- | --- | --- | --- |
| 1 | Strict Clock | Constant Size | Brownian | -7460.8 | -7460.8 |
| 2 | UCLN | Constant Size | Brownian | -7449.3 | -7449.0 |
|  | UCLN | GMRF Bayesian Skyride | Brownian | -7451.9 | -7451.3 |
| 3 | UCLN | Exponential | Brownian | -7446.2 | -7446.2 |
| *4* | UCLN | Exponential | Lognormal RRW | -7413.7 | 7413.8 |
|  | *UCLN* | *Exponential* | *Cauchy RRW* | *-7383.5* | *-7383.6* |

UCLN: Uncorrelated relaxed clock with a log normal distribution.
